# genomesizeR: An R package for genome size prediction

**DOI:** 10.1101/2024.09.08.611926

**Authors:** Celine Mercier, Joane Elleouet, Loretta Garrett, Steve A Wakelin

## Abstract

The genome size of organisms present in an environment can provide many insights into evolutionary and ecological processes at play in that environment. The genomic revolution has enabled a rapid expansion of our knowledge of genomes in many living organisms, and most of that knowledge is classified and readily available in the databases of the National Center for Biotechnology Information (NCBI). The `genomesizeR` tool leverages the wealth of taxonomic and genomic information present in NCBI databases to infer the genome size of Archeae, Bacteria, or Eukaryote organisms identified at any taxonomic level. This R package uses statistical modelling on data from the most up-to-date NCBI databases and provides three statistical methods for genome size prediction of a given taxon, or group of taxa. A straightforward ‘weighted mean’ method identifies the closest taxa with available genome size information in the taxonomic tree, and averages their genome sizes using weights based on taxonomic distance. A frequentist random effect model uses nested genus and family information to output genome size estimates. Finally a third option provides predictions from a distributional Bayesian multilevel model which uses taxonomic information from genus all the way to superkingdom, therefore providing estimates and uncertainty bounds even for under-represented taxa.

All three methods use:

- A list of queries; a query being a taxon or a list of several taxa. The package was designed to make it easy to use with data coming from environmental DNA experiments, but works with any table of taxa.
- A reference database containing all the known genome sizes, built from the NCBI databases, with associated taxa, provided in an archive to download.
- A taxonomic tree structure as built by the NCBI, provided in the same archive.

`genomesizeR` retrieves the taxonomic classification of input queries, estimates the genome size of each query, and provides 95% confidence intervals for each estimate.

## Statement of need

The size of microbial genomes and its evolution can provide important insights into evolutionary and ecological processes influencing both microbial species and the environments in which they inhabit. The shedding of unnecessary genetic elements and their associated biosynthetic pathways, for example, is a common phenomenon observed in organisms with a high degree of host symbiosis (Moran 2002; Brader et al. 2014; Vandenkoornhuyse et al. 2007). Genome size reduction has also been observed in organisms experiencing arid environments (Liu et al. 2023), or a narrow range of substrates or metabolic options (Tyson et al. 2004). Among many others, these findings demonstrate the opportunities associated with including genome size as a key trait in microbial communities to provide insights spanning niche size, co-evolution, adaption, and metabolic flexibility of the microbiomes present, but also stability, and ecophysiological and functional complexity of abiotic and biotic environments.

However, characterizing genome size for all organisms in a microbiome remains challenging. The exponentially growing genome databases are an inexpensive resource unlocking a myriad of research opportunities in all fields of environmental sciences, but genome size estimates for many taxa found in environmental samples are missing from public databases, or fully unknown. The evolutionary rule that phylogenetically related organisms share genetic similarities can be exploited and genome size for taxa with unknown genome size can be statistically inferred from related taxa with known genome size, using taxonomy as a proxy for phylogeny. Another challenge is the precision of identification: some taxa can only be identified at high taxonomic levels. Statistical methods can also be used to infer their genome size range from databases. To our knowledge, there is no convenient and fast way to obtain genome size estimates with uncertainty bounds for all organisms identified or partially identified in an environmental sample.

Using the increased prevalence of whole-genome information for all organisms, we have therefore developed `genomesizeR`, allowing the inference of genome size of many queries at once, based on taxonomic information and available genome data from the NCBI.

## Methods

### NCBI database filtering and processing

The reference database used is built by querying all genome metadata information from the curated NCBI RefSeq database (O’Leary et al. 2016). Filters are applied to only keep full genomes, and discard data that the NCBI has tagged as anomalous, or as abnormally large or small sizes.

This raw database is then prepared to include more pre-computed information to be used by the package. Genome sizes are aggregated to the species level by iteratively averaging all entries below, and estimated standard errors for each species with multiple genome size entries are stored. The package can therefore only provide estimates at the level of species and above. Average genome sizes and their associated standard error values are also pre-computed, to be used by the weighted mean method.

### Bayesian method

The NCBI database of species with known genome sizes was split by superkingdom (Bacteria, Archeae, Eukaryotes). A distributional Bayesian linear hierarchical model using the `brm` function from the `brms` package (Bürkner 2017) was fitted to each superkingdom dataset. The general model structure is outlined below.

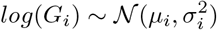

where *G*_*i*_ is the genome size of species *i* in the units of 10 Mbp. The model uses predictors for both mean and standard deviation. The mean is modelled as follows:

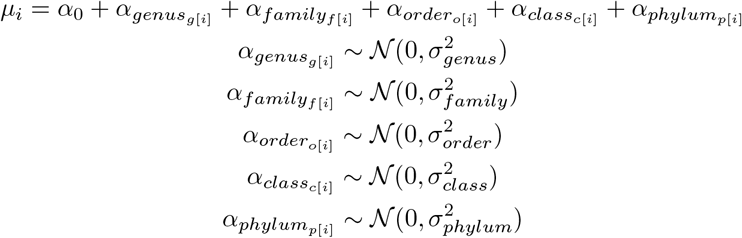

The following prior distributions are used:

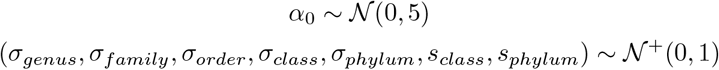

Differences in genome size variability was observed among taxa, therefore the model also adds predictors to the standard deviation of the response. The standard deviation is modelled as follows:

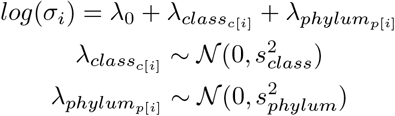

with priors

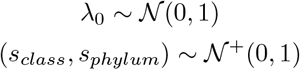

𝒩^+^ is the normal distribution truncated to positive values. *g*[*i*],*f*[*i*],*o*[*i*],*c*[*i*] and *p*[*i*] are respectively the index for the genus, family, order, class, and phylum of entry *i* in the species-level database. Note that taxonomic groups are naturally nested within each other and the indices are designed to be unique to the particular taxonomic group it represents.

The estimation process uses Stan’s Hamiltonian Monte Carlo algorithm with the U-turn sampler.

Posterior predictions are obtained using the `predict` function from the `brms` package, and 95% credible intervals are obtained using 2.5% and 97.5% quantiles from the posterior distribution.

Queries corresponding to identified species with an available genome size estimate in the NCBI database get allocated the genome size value of the database (averaged at the species level) and 95% confidence intervals are calculated based on the standard error of the mean of all genome sizes available for that species in the processed NCBI database.

### Frequentist method

A frequentist linear mixed-effects model using the `lmer` function from the `lme4` package (Bates et al. 2015) was fitted to the NCBI database of species with known genome sizes. The model is as follows:

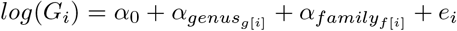

where *α*_0_ is the overall mean, 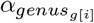 and 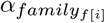 are random effect of genus and family for genus *g*[*i*] and family *f* [*i*] and *e*_*i*_ is the residual error of observation *i*.

The estimation process using the restricted maximum likelihood method (REML). A prediction interval is computed using the `predictInterval` function from the `merTools` package (Knowles and Frederick 2024). As higher nested levels (order, class) are not taken into account in the model, predictions produced for queries above the family level are not to be trusted.

Queries corresponding to identified species with an available genome size estimate in the NCBI database get allocated the genome size value of the database (averaged at the species level) and 95% confidence intervals are calculated based on the standard error of the mean of all genome sizes available for that species in the processed NCBI database.

### Weighted mean method

The weighted mean method computes the genome size of a query by averaging the known genome sizes of surrounding taxa in the taxonomic tree, with a weighted system where further neighbours have less weight in the computed mean. The identification of related taxa is limited to levels below and including order.

The pseudocode describing the algorithm for the estimate and confidence interval computation is available in the package vignettes.

For queries relating to well-characterised species where many genetic studies have been performed, such as model organisms, this might lead to more precise predictions than the two other methods. This method can also perform better than the others if your queries consist of lists of taxa (for example, an output of *blastn* where several matches can be obtained for each query). Otherwise, we suggest using one of the other methods, as the confidence intervals calculated are less reliable for the weighted mean method.

### Implementation

The main steps of all methods are multithreaded on POSIX systems using the packages parallel (R Core Team 2024) and doParallel (Corporation and Weston 2022).

The packages ncbitax (Machne 2022) and CHNOSZ (Dick 2019) are used to read the taxonomy data, dplyr (Wickham et al. 2023) and biomformat (McMurdie and Paulson 2024) are used for some of the formatting, and pbapply (Solymos and Zawadzki 2023) is used to display the progress bar.

The R package accepts as input formats the common ‘taxonomy table’ format used by popular packages such as phyloseq (McMurdie and Holmes 2013) and mothur (Schloss et al. 2009), and any file or data frame with a column containing either NCBI taxids or taxon names. The output format is a data frame with the same columns as the input, with some added columns providing information about the estimation and the quality of the estimation. The user can also choose a simple output format only containing the estimation information.

Several plotting functions using the ggplot2 (Wickham 2016) and ggtree (Yu 2020) packages are also provided to visualise the results.

### Example

This example data is a subset of the dataset from Labouyrie et al. (2023). First, the genome sizes are predicted from the taxa:

~~~
results = estimate_genome_size(example_input_file, refdata_archive_path, sep=‘\t’,
match_column=‘TAXID’, output_format=‘input’, ci_threshold = 0.3)
#############################################################################
# Genome size estimation summary:
#
#  22.22222 % estimations achieving required precision
#
   Min.   1st Qu.    Median     Mean   3rd Qu.       Max.
3007721  5408472  16980834  23969767  41811396  143278734
# Estimation status:
Confidence interval to estimated size ratio > ci_threshold     OK
                                                    140      40
~~~

Then, the results can be visualized using the plotting functions provided. Figure 1 shows a histogram and a boxplot of the estimated genome sizes for each sample. Figure 2 shows a tree showing the taxonomic relationships as well as the estimated genome sizes. The difference between the genome size distribution of bacteria (16S marker) and fungi (ITS marker) is visible in both figures, as well as different distributions between samples.

**Figure 1:**
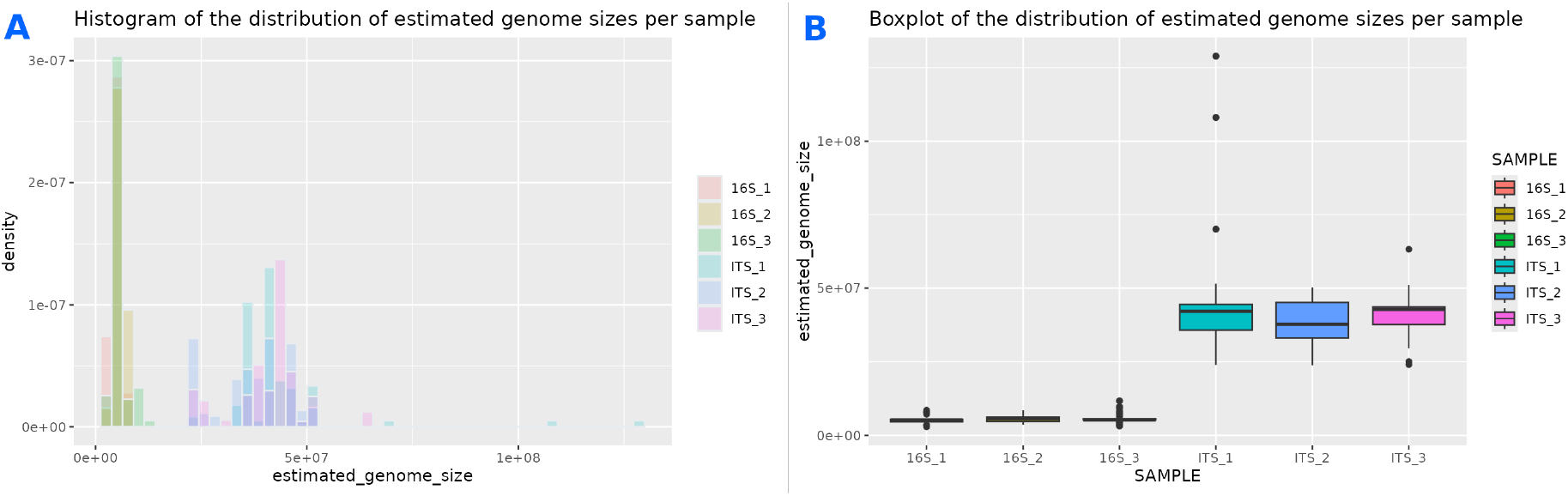
Histogram (A) and boxplot (B) of estimated genome sizes for each sample

**Figure 2:**
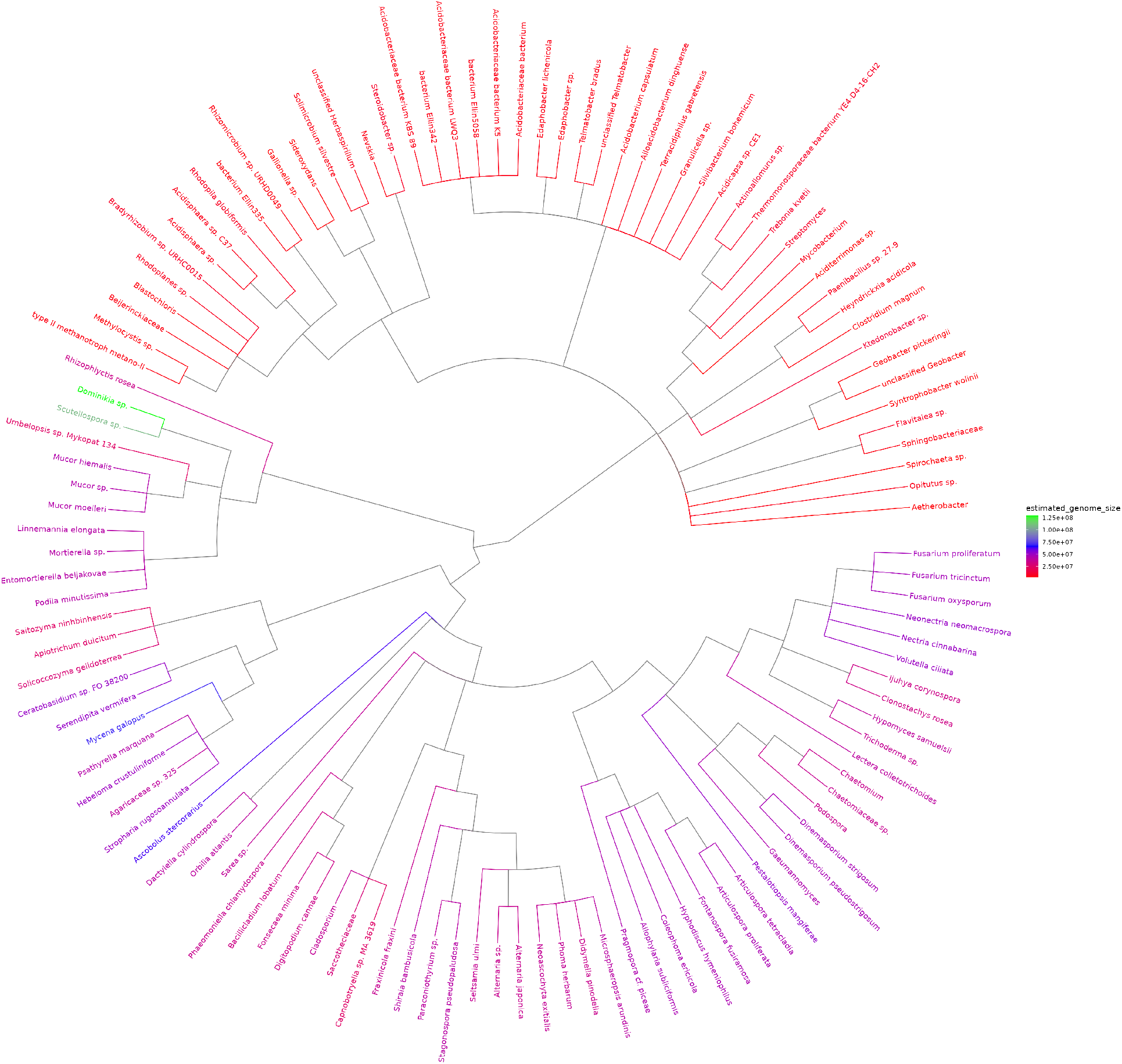
Tree representing taxonomic relationships and estimated genome sizes between queries

### Method comparison

The applicability of each method varies (Table 1). The Bayesian method outputs results for any taxon that is recognised in the NCBI taxonomy. The frequentist random effects model method only outputs results for queries that have a match at the species, genus, or family level. The weighted mean method only performs an estimation for queries that have at least two matches at the species, genus, family, or order level.

**Table 1:**
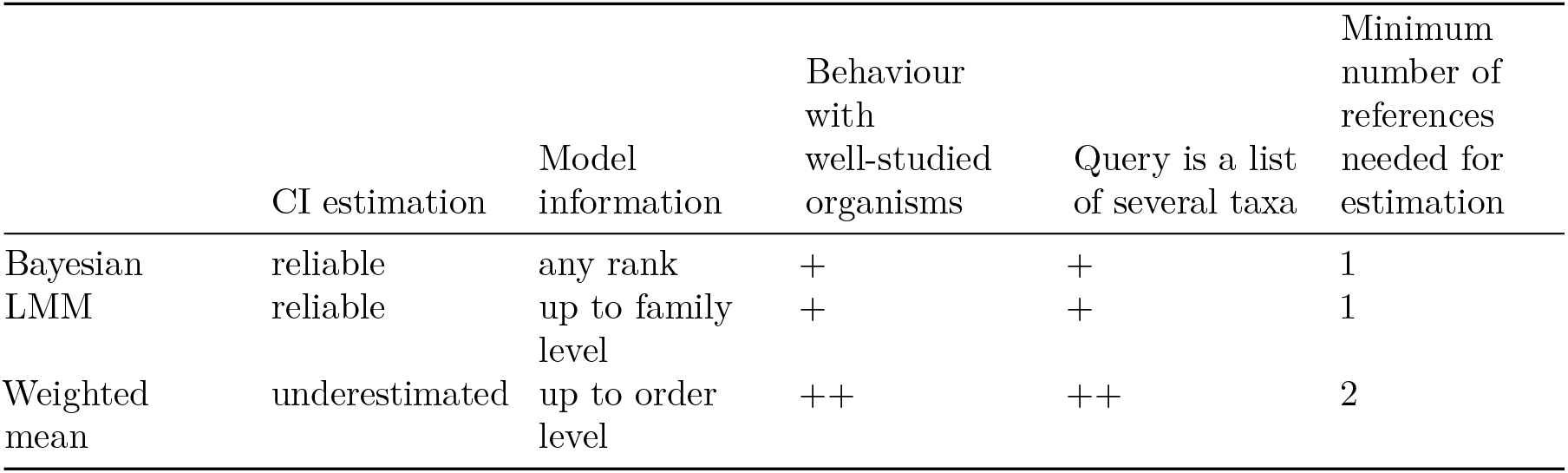
Comparison of method behaviour and applicability.

|  | CI estimation | Model information | Behaviour with well-studied organisms | Query is a list of several taxa | Minimum number of references needed for estimation |
| --- | --- | --- | --- | --- | --- |
| Bayesian | reliable | any rank | + | + | 1 |
| LMM | reliable | up to family level | + | + | 1 |
| Weighted mean | underestimated | up to order level | ++ | ++ | 2 |

Below is a comparison of estimates for an example set of bacteria and fungi queries where the highest level of match with the database is the family level. Estimates and the width of confidence intervals differ between methods (figures Figure 3 and Figure 4).

**Figure 3:**
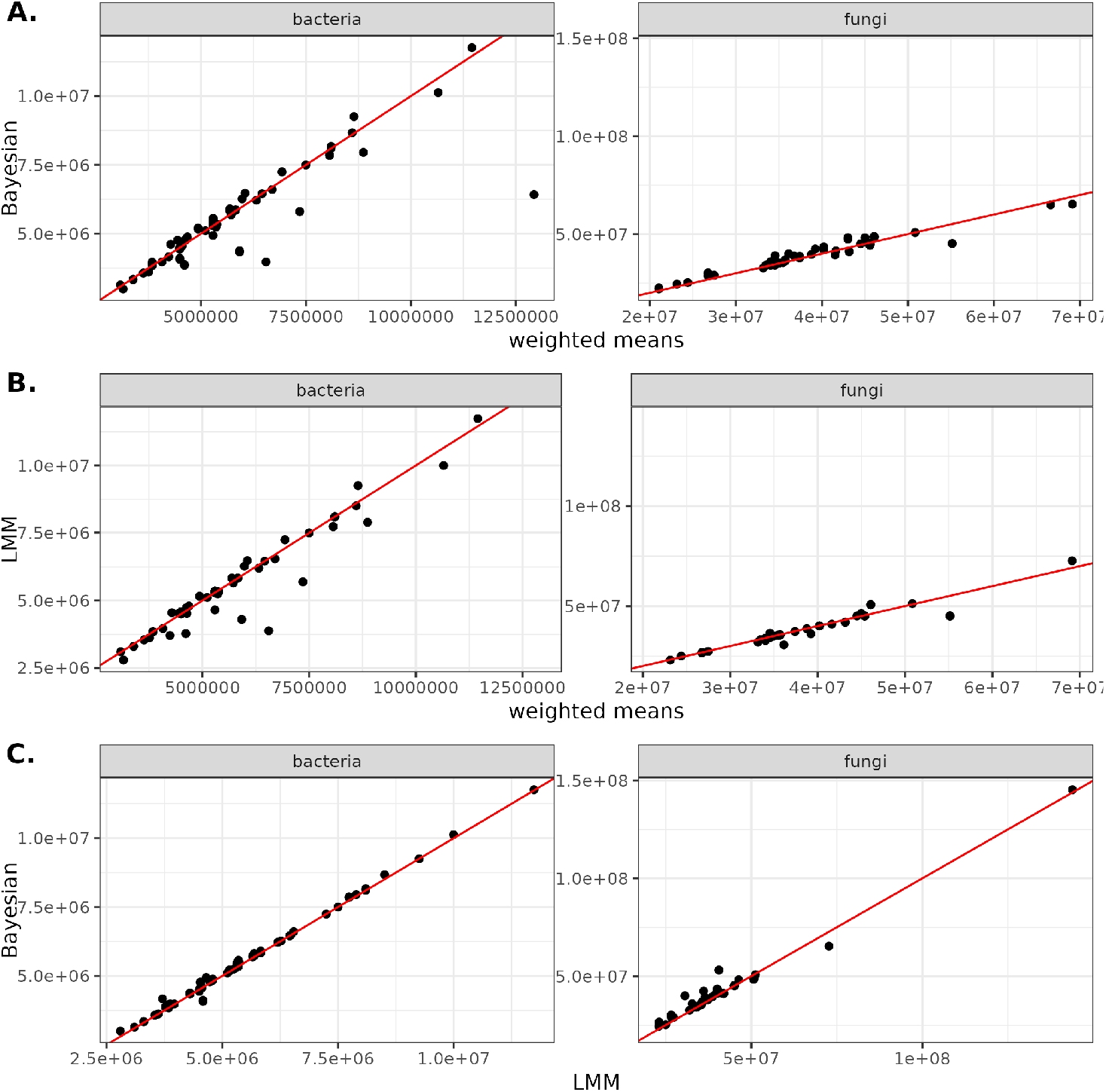
Pairwise comparison of estimates from different methods for bacteria and fungi. A. Bayesian model vs. weighted means method; B. Frequentist mixed model vs. weighted means method; C. Bayesian model vs. frequentist mixed model.

**Figure 4:**
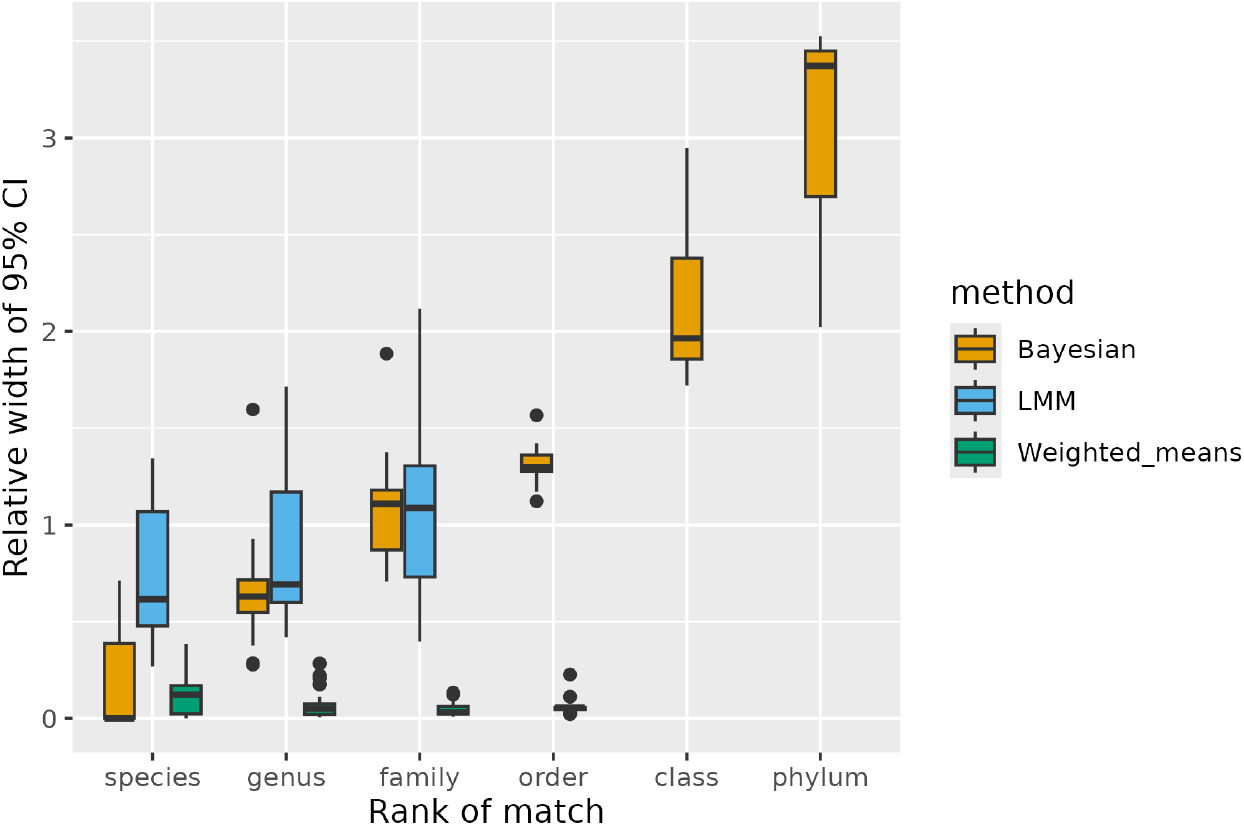
Distribution of relative 95% confidence intervals per method and rank match. 95% confidence intervals were scaled by estimated size. Rank match refers to the smallest rank in common between the query and the closest database entry with a valid genome size.

## Availability

- Project name: genomesizeR
- Project home page: https://github.com/ScionResearch/genomesizeR
- Operating system(s): Platform independent
- Programming language: R
- License: GNU General Public License

## Acknowledgements

We acknowledge contributions from Sean Husheer. The authors declare that they have no conflict of interest. Funding for this research came from the Tree-Root-Microbiome programme, which is funded by MBIE’s Endeavour Fund and in part by the New Zealand Forest Growers Levy Trust (C04X2002). We make no warranties regarding the accuracy or integrity of the Data. We accept no liability for any direct, indirect, special, consequential or other losses or damages of whatsoever kind arising out of access to, or the use of the Data. We are in no way to be held responsible for the use that you put the Data to. You rely on the Data entirely at your own risk.

## Notes

https://github.com/ScionResearch/genomesizeR

https://doi.org/10.5281/zenodo.13733184

### Competing Interest Statement

The authors have declared no competing interest.

